# Dopamine transporter trafficking and Rit2 GTPase: Mechanism of action and *in vivo* impact

**DOI:** 10.1101/712448

**Authors:** Rita R. Fagan, Patrick J. Kearney, Carolyn G. Sweeney, Dino Luethi, Florianne E. Schoot Uiterkamp, Klaus Schicker, Brian S. Alejandro, Lauren C. O’Connor, Harald H. Sitte, Haley E. Melikian

## Abstract

Following its evoked release, DA signaling is rapidly terminated by presynaptic reuptake, mediated by the cocaine-sensitive DAT. DAT surface availability is dynamically regulated by endocytic trafficking, and direct PKC activation acutely diminishes DAT surface expression by accelerating DAT internalization. Previous cell line studies demonstrated that PKC-stimulated DAT endocytosis requires both Ack1 inactivation, which releases a DAT-specific endocytic brake, and the neuronal GTPase, Rit2, which binds DAT. However, it is unknown whether Rit2 is required for PKC-stimulated DAT endocytosis in DAergic terminals, or whether there are region- and/or sex-dependent differences in PKC-stimulated DAT trafficking. Moreover, the mechanisms by which Rit2 controls PKC-stimulated DAT endocytosis are unknown. Here, we directly examined these important questions. *Ex vivo* studies revealed that PKC activation acutely decreased DAT surface expression selectively in ventral, but not dorsal, striatum. AAV-mediated, conditional Rit2 knockdown in DAergic neurons impacted baseline DAT surface:intracellular distribution in DAergic terminals from female ventral, but not dorsal, striatum. Further, Rit2 was required for PKC-stimulated DAT internalization in both male and female ventral striatum. FRET and surface pulldown studies in cell lines revealed that PKC activation drives DAT-Rit2 surface dissociation, and that the DAT N-terminus is required for both PKC-mediated DAT-Rit2 dissociation and DAT internalization. Finally, we found that Rit2 and Ack1 independently converge on DAT to facilitate PKC-stimulated DAT endocytosis. Together, our data provide greater insight into mechanisms that mediate PKC-regulated DAT internalization, and reveal unexpected region-specific differences in PKC-stimulated DAT trafficking in *bona fide* DAergic terminals.

---

DA neurotransmission is required for motor control, learning, memory, motivation, and reward (1,2). DAergic dysregulation is evidenced in numerous neuropsychiatric disorders, including ADHD, ASD, schizophrenia, bipolar disorder, addiction, and PD (3-8). DA signaling is tightly controlled by the presynaptic DAT, which rapidly clears synaptically released DA. DAT is also the primary target for addictive and therapeutic psychostimulants, including AMPH, cocaine, and methylphenidate (Ritalin), which inhibit DAT as competitive substrates (AMPH) and antagonists (cocaine and methylphenidate) (9). Genetic DAT deletions in mice and *Drosophila melanogaster* elevate extracellular DA concentrations and evoke hyperactivity (10-12), and human DAT missense mutations have been reported in PD, ADHD and ASD patients (13-19). Together, these studies underscore that DAT is critical to maintain DAergic homeostasis (9).

Given its central role in DAergic signaling, intrinsic neuronal mechanisms that alter DAT surface expression and function are likely to significantly impact DAergic transmission. DAT surface availability is dynamically modulated by endocytic trafficking (20-22). A negative regulatory mechanism, or “endocytic brake”, tempers basal DAT endocytosis (23,24), and acute PKC activation disengages the DAT endocytic brake, stimulates DAT internalization, and rapidly diminishes DAT surface expression (25,26). The DAT N- and C-termini encode residues required to engage the DAT endocytic brake, and, when mutated, markedly accelerate DAT internalization (15,23,27). We previously reported that the non-receptor tyrosine kinase, Ack1 (AKA: TNK2), is a critical component of the DAT endocytic brake, and that Ack1 inactivation is required for PKC-mediated brake release (24).

Rit2 (AKA: Rin) is a neuronal small GTPase that lacks a CAAX domain, and associates with the plasma membrane in a phosphoinositide-dependent manner (28-30). Rit2 gene expression is highly enriched in DA neurons (31), and several recent GWAS studies identified Rit2 SNPs and long tandem repeat variants associated with multiple DA-related disorders, including PD, ASD, and schizophrenia (32-37). However, despite its disease association, relatively little is known about endogenous DAergic Rit2 function. We previously reported that DAT directly binds to Rit2, and that Rit2 activity is required for PKC-stimulated DAT internalization (38). Moreover, we recently found that *in vivo* DAergic Rit2 knockdown (Rit2-KD) differentially alters acute cocaine sensitivity in males and females (39). However, it remains unknown whether PKC-stimulated DAT internalization in DAergic terminals requires Rit2, and whether there are region- or sex-specific differences in DAT’s reliance upon Rit2. Further, it is unclear how intrinsic DAT domains influence the DAT-Rit2 interaction, or whether Rit2 and Ack1 coordinate to release the DAT endocytic brake. In the present study, we leveraged biochemical and genetic approaches, in both cultured cells and *ex vivo* mouse striatal slices, to directly address these salient questions.

## RESULTS

### Rit2 cellular expression and antibody specificity

In our previous study, in which we initially reported the DAT-Rit2 (Rin) interaction (38), there were several paradoxical findings regarding 1) how PKC activation impacted the DAT-Rit2 interaction, and 2) the Rit2 expression profile across various cell lines. In cellular imaging studies, which used CFP-Rit2, Rit2 appeared to remain at the plasma membrane following PKC-stimulated DAT internalization, suggesting that PKC may drive DAT and Rit2 to dissociate. In contrast, parallel co-IP studies found that PKC activation <u>increased</u> the DAT-Rit2 interaction in PC12 cells. Furthermore, although Rit2 expression is reportedly restricted to neurons (28,29,31,40), we detected a single, ∼20kDa immunoreactive band by immunoblot in all neuronal and non-neuronal cell lines tested, as well as Rit2 mRNA expression in these cell lines by standard RT-PCR. The previous studies utilized the anti-Rit2 monoclonal antibody, clone 27G2, and in that report we additionally confirmed that 27G2 specifically recognizes Rit2, but not Rit1, (the ubiquitously expressed Rit2 homologue) using fluorescently-tagged Rit2 and Rit1 fusion proteins. Recently, we decided to take advantage of highly-specific, Rit2-directed, real-time qPCR probes to re-assess the Rit2 expression profile across a large panel of human, mouse and rat cell lines, as well as in mouse and rat midbrain, all of which exhibit the single, 20kDa immunoreactive band when probed with the anti-Rit2 27G2. To our surprise, Rit2 mRNA was undetectable in any of the mouse or rat cell lines tested, whereas a robust Rit2 mRNA signal was detected in both mouse and rat midbrain controls (Table I). Moreover, among the human cell lines tested, Rit2 mRNA was only detected in SK-N-DZ cells, as previously reported (40), and at low levels in SH-SY5Y cells. However, SK-N-MC and HEK293T cells expressed markedly less/negligible Rit2 signal than SK-N-DZ cells, ranging from 20-300-fold less (Table I).

Our current RT-qPCR results raised the possibility that although 27G2 recognizes Rit2, it also may cross-react with a ubiquitously expressed protein that has an electrophoretic mobility close to that of Rit2. To test this, we screened several newer, commercially available anti-Rit2 antibodies using cell lysates from HEK293T cells transfected with CFP-Rit2. Consistent with our previous report, clone 27G2 identified a single 20kDa band in both transfected and non-transfected cells, and also detected CFP-Rit2 selectively in transfected cells (Fig. 1A). However, when immunoblots were probed with αRit2 clone 4B5, the low molecular weight immunoreactive band was not detected, while CFP-Rit2 was detected in transfected cells (Fig. 1A). These results are consistent with our RT-qPCR results and confirm that 27G2 detects an artifactual band with an electrophoretic mobility close to the Rit2 predicted size. We next used 4B5 to assess Rit2 in cells transfected with HA-Rit2, and in mouse DAergic tissues. Clone 4B5 detected a single, ∼28kDa immunoreactive band selectively in cells transfected with HA-Rit2, which was identical in mobility to an immunoreactive band detected with αHA, in parallel (Fig. 1B). αHA also detected a higher molecular weight band (∼30kDa) specifically in cells transfected with HA-Rit2, which we did not detect using 4B5 (Fig. 1B), suggesting that Rit2 may have multiple isoforms, and that 4B5 may only detect one of these. 4B5 also detected bands at ∼24kDa in mouse lysates enriched for either dorsal striatum or ventral midbrain (Fig. 1B). These immunoblot results were consistent with our RT-qPCR results, and confirmed that most cell lines either do not express Rit2, or express Rit2 at negligible levels. They further support that the 27G2 antibody cannot reliably distinguish between Rit2 and a robust, artifactual, background band. It should also be noted that while 4B5 specifically detects Rit2, it does so with extremely low sensitivity, even when Rit2 is highly overexpressed. Indeed, using the 4B5 antibody, we could only detect Rit2 by immunoblot in lysates from mouse tissues when a very high protein mass was loaded. Moreover, although both hRit2 and mRit2 are predicted to have identical number of amino acids, and a predicted mass of ∼24kDa each, the overexpressed hRit2 protein ran slightly higher than the putative mRit2 band. There are two known hRit2 isoforms, variants 1 and 2, which are predicted to be ∼24kDa and ∼17kDa, respectively, and our hRit2 cDNA codes for variant #1. To date it is unknown whether there are tissue specific Rit2 isoforms in mouse neurons. Given that there is no global Rit2^-/-^ mouse available, we therefore cannot say with absolute certainty that the single immunoreactive band in mouse tissue is definitively Rit2, or possibly a smaller splice variant. Therefore, for the majority of our cell line studies, we opted to use HA-Rit2 for greater sensitivity in cell lines and did not further assess Rit2 protein in tissue.

**Figure 1.**
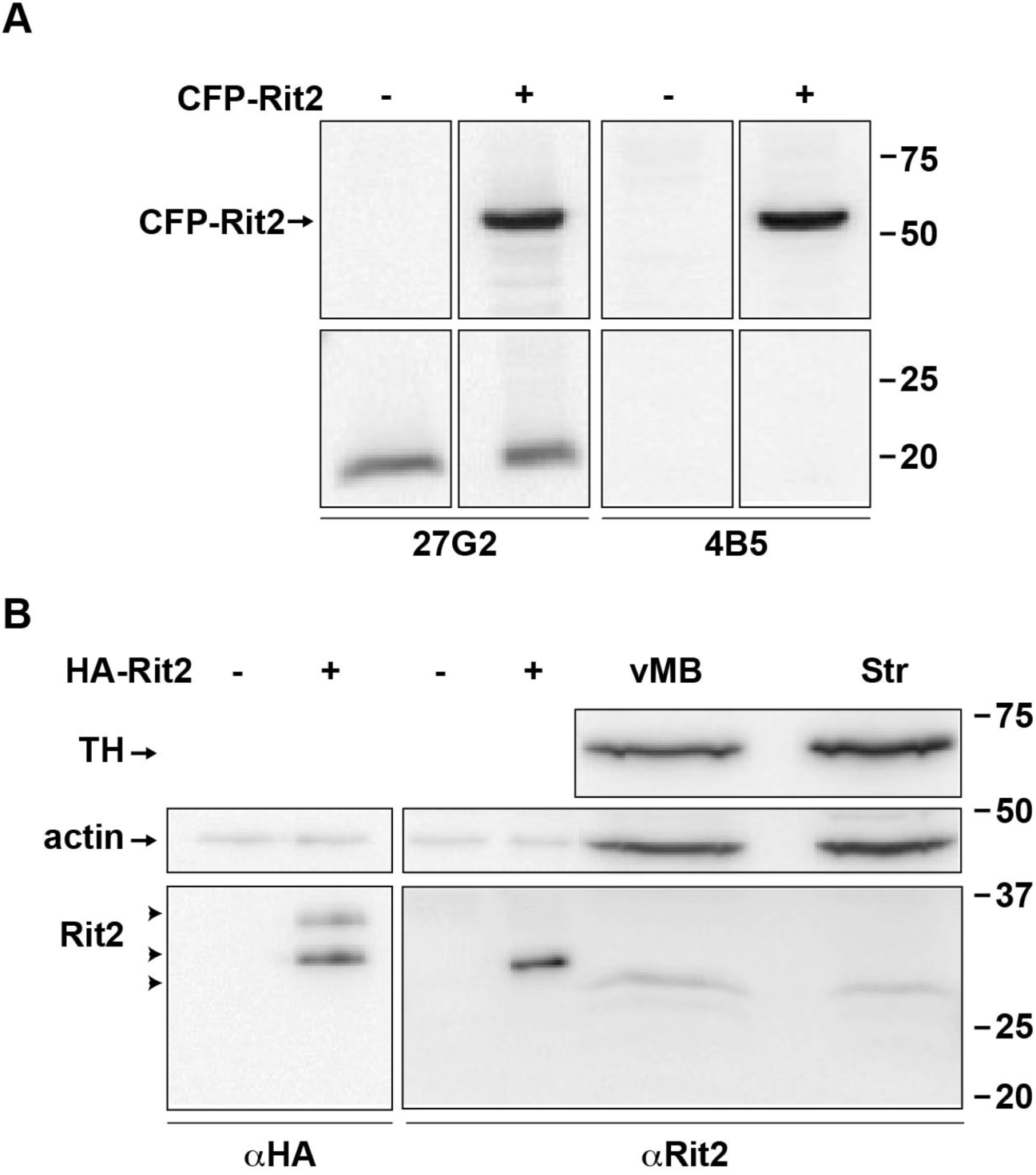
Rit2 protein is specifically detected by clone 4B5, but not clone 27G2, *α*Rit2 antibodies. *Rit2 detection in transfected cell lines and mouse tissues by immunoblot.* **A.** HEK293T cells were transfected with either vector (-) or CFP-Rit2 (+) and cell lysates were assessed by immunoblot 48 hrs. post-transfection, probing with either αRit2 clones 27G2 (left) or 4B5 (right) as described in *Materials and Methods*. Molecular weight markers indicate kDa. **B.** HEK293T cells transfected with vector (-) or HA-Rit2 (+), mouse ventral midbrain (vMB), and striatum (Str), were assessed by immunoblot, probing with either αHA (left) or αRit2 clone 4B5 (right). 10µg/lane and 100µg/lane were loaded for transfected cell lysates and mouse brain lysates, respectively. Arrowheads indicate Rit2 immunoreactive bands.

### Rit2 is required for PKC-stimulated, but not basal, DAT internalization

In our original DAT-Rit2 study (38), we used shRNA to silence Rit2 in SK-N-MC cells and to test whether Rit2 is required for PKC-stimulated DAT downregulation. In light of our current findings that SK-N-MC cells do not appreciably express Rit2, we were prompted to 1) re-screen Rit2 targeted shRNAs, and 2) reassess whether Rit2 is required for PKC-stimulated DAT internalization in SK-N-DZ cells, which endogenously express Rit2. We screened several candidate human Rit2-directed shRNAs, and identified two shRNAs (shRit2-104 and −107) that significantly silenced CFP-hRit2 protein expressed in Neuro2a cells (Fig. 2A). Additionally, both shRit2-104 and −107 significantly silenced endogenous Rit2 mRNA expression in SK-N-DZ cells (Fig. 2B). We utilized shRit2-107 to ask whether Rit2 is required PKC-stimulated DAT internalization in SK-N-DZ cells. Rit2-KD significantly blocked PKC-stimulated DAT endocytosis as compared to vector-transduced cells (Fig. 2C), consistent with a requisite role for Rit2 in PKC-stimulated DAT internalization. The ability of shRit2-107 to block PKC-stimulated DAT endocytosis was not likely due to off-target effects, as shRit2-107 did not decrease expression of Rit1, the closest homolog to Rit2 (Fig. 2D). To further assure that shRit2-107 effects were specific, we repeated these studies with shRit2-104. Similar to our findings with shRit2-107, Rit2 silencing with shRit2-104 significantly blocked PKC-stimulated DAT internalization (Fig. 2E).

**Figure 2.**
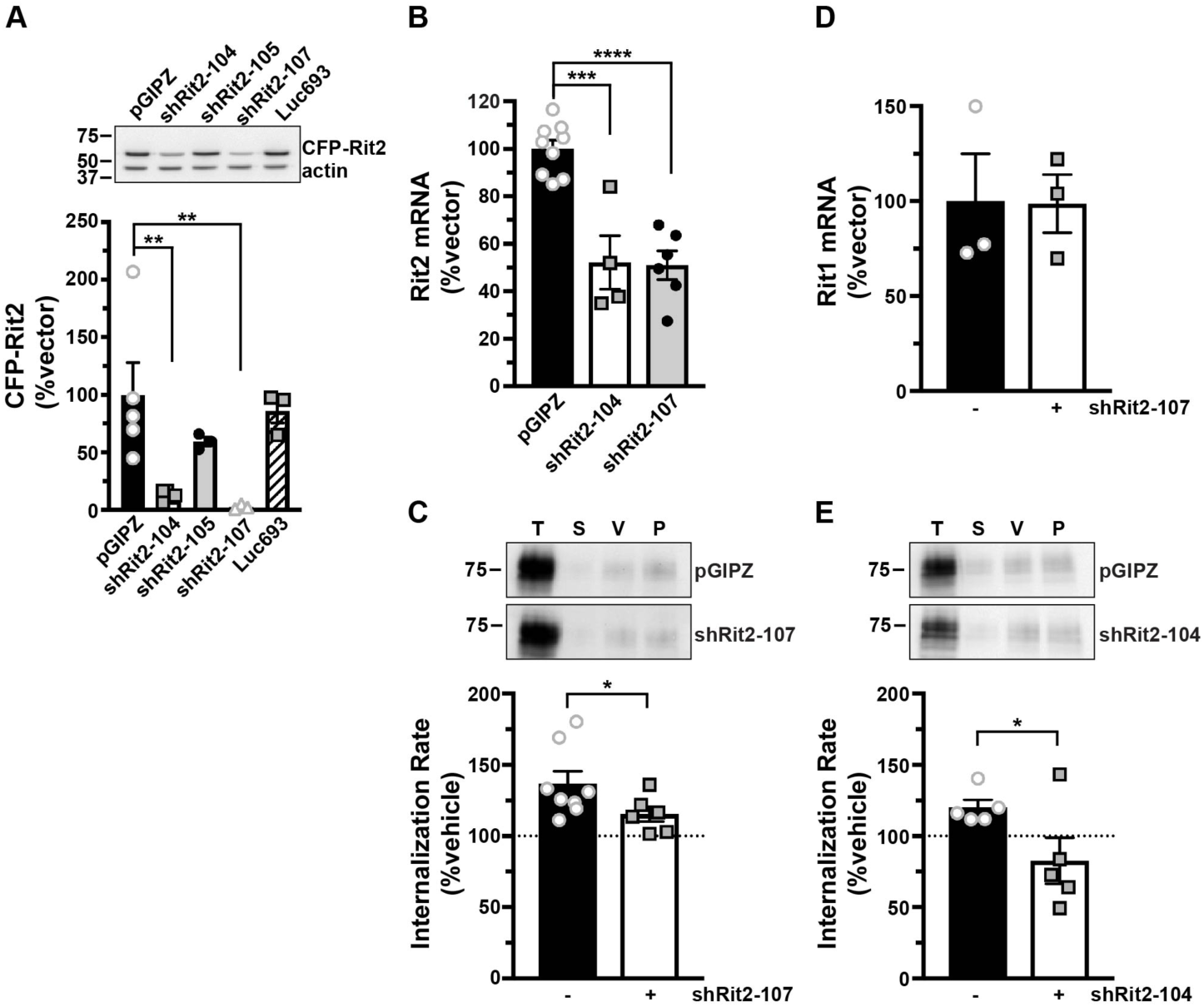
Rit2 is required for PKC-stimulated DAT internalization in DAT-SK-N-DZ cells. **A.** *Human Rit2 shRNA screen*. Mouse N2a cells were co-transfected with human CFP-Rit2, and either pGIPZ vector, luciferase293 control, or the indicated Rit2 shRNA-pGIPZ plasmids. *Top*: Representative immunoblots. *Bottom*: Average CFP-Rit2 levels, normalized to actin loading controls, expressed as %pGIPZ vector ±S.E.M. Asterisks indicate a significant difference from pGIPZ controls Asterisks indicate a significant difference from pGIPZ controls (one-way ANOVA F_(4, 14)_ = 6.396, p = 0.004; Dunnett’s multiple comparisons test, pGIPZ vs. shRit2-104: **p = 0.005; vs. shRit2-105: p = 0.36; vs shRit2-107: **p = 0.006; vs luc693: p = 0.95, n=3-5). **B.** *shRit2-mediated knockdown in stable DAT-SK-N-DZ cells*. DAT-SK-N-DZ cells were transduced with control, shRit2-104, or shRit2-107 lentiviral particles, and Rit2 mRNA expression was measured by RT-qPCR 96 hrs post-transduction. Average □ □Ct values are presented, expressed as %control-transduced values ±S.E.M. Asterisks indicate a significant difference from pGIPZ controls (one-way ANOVA F_(2, 16)_ = 25.09, p<0.0001; Dunnett’s multiple comparisons test: pGIPZ vs shRit2-104: ***p = 0.0001; vs shRit2-107: ****p<0.0001, n=4-9). **C.** *DAT internalization assay*. Stable DAT-SK-N-DZ cells were transduced with the indicated lentiviral particles and DAT internalization rates were measured 96 hrs post-transduction as described in *Materials and Methods. Top*: Representative immunoblots depicting surface DAT expression at t=0 (T), strip control (S), and internalized DAT during either vehicle (V) or 1 µM PMA (P) treatment. *Bottom*: Average DAT internalization rates expressed as %vehicle-treated control rates ±S.E.M. *p=0.04, one-tailed Student’s t test, n=6-8. **D.** *Rit1 mRNA expression specificity control*. DAT-SK-N-DZ cells were transduced with control, or shRit2-107 lentiviral particles, and Rit1 expression was measured by RT-qPCR 96 hrs post-transduction. Average ΔΔCt values are presented, expressed as %control-transduced values ±S.E.M. shRit2-107 transduction did not significantly affect Rit1 mRNA expression, p=0.965, two-tailed Student’s t test, n=3**. E.** *DAT internalization assays*. Stable DAT-SK-N-DZ cells were transduced with the indicated lentiviral particles and DAT internalization rates were measured 96 hrs post-transduction as described in *Materials and Methods*. Data are presented identically as in (C). *p=0.03, one-tailed Student’s t test, n=5.

### Rit2 is required for striatal steady state DAT surface expression and PKC-stimulated DAT internalization in a region- and sex-specific manner

PKC-stimulated DAT internalization in response to phorbol ester treatment has been reported by numerous laboratories, in a variety of transfected cell lines (15,25,41-45). Moreover, we previously reported that phorbol ester (PMA) treatment decreases DAT surface levels in DAergic terminals in *ex vivo* total striatal slices containing both dorsal (DS) and ventral (VS) striatum (26). However, it is unknown whether PKC-stimulated DAT internalization differs between DAergic terminal regions, such as DS and VS, or between males and females. Moreover, although Rit2 is required for PKC-stimulated DAT internalization in SK-N-DZ cells (Fig. 2), it is unknown whether Rit2 is required for PKC-stimulated DAT endocytosis in DAergic terminals. We recently leveraged the TET-OFF system to achieve conditional, inducible DAergic Rit2-KD in *Pitx3*^*IRES2-tTA*^ mice, in which AAV9-shRit2 injection into mouse VTA significantly silenced Rit2 expression in both VTA and SNc (39). We found that DAergic Rit2 silencing decreased total striatal DAT protein in males, but not females. Further, although total DAT protein decreased within male total striatum, the DAT surface:intracellular ratio was unchanged in either DS or VS, resulting in less overall surface DAT in both male striatal subregions (39). In the current study, we extended our *in vivo* Rit2-KD studies and asked whether Rit2 impacts DAT basal distribution in female DS and VS. We further asked whether PKC-mediated DAT internalization requires Rit2 in male and female DS and VS. Male and female *Pitx3*^*IRES2-tTA/+*^ mice VTA were bilaterally injected with either AAV9-TRE-eGFP or AAV9-TRE-shRit2, and DAT surface expression was measured by *ex vivo* slice biotinylation in the VS and DS, following treatment ±1µM PMA, 30 min, 37°C.

We first examined the effect of Rit2 silencing on DAT surface expression in female DS and VS, under both basal and PKC-stimulated conditions. Surprisingly, in DS, PKC activation did not decrease DAT surface expression in control female mice (Fig. 3A). Moreover, DAergic Rit2-KD had no effect on DS DAT surface expression, under either basal or PKC-stimulated conditions (Fig. 3A). In contrast, in female VS, PKC activation significantly reduced DAT surface levels, and DAergic Rit2-KD completely blocked further surface loss in response to PKC activation (Fig. 3B). Moreover, DAergic Rit2-KD significantly reduced the basal DAT surface level in female VS, as compared to control mice (Fig. 3B).

**Figure 3.**
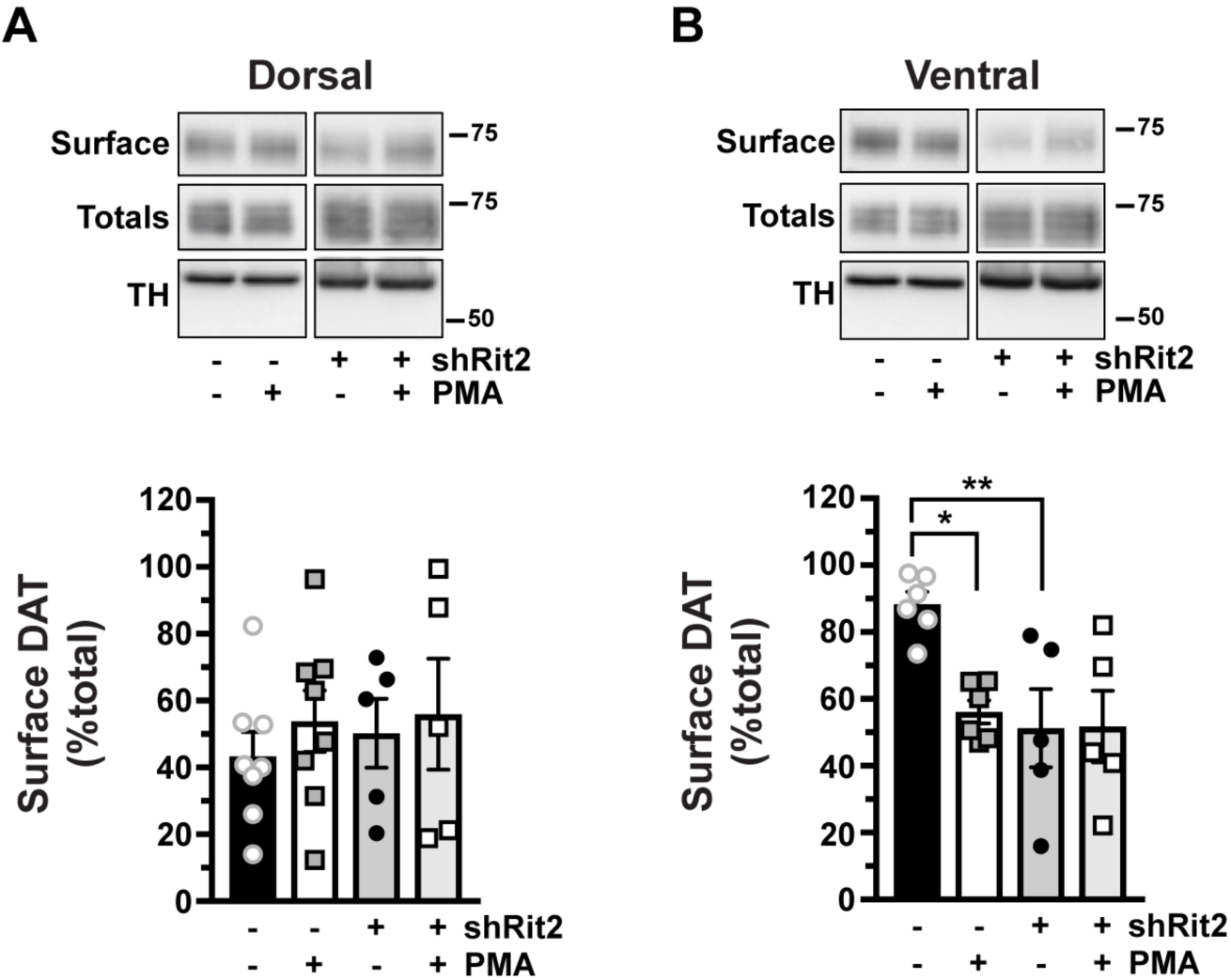
PKC-induced DAT internalization in females is limited to ventral striatum and requires Rit2. *Conditional Rit2 silencing in DA neurons and ex vivo striatal slice biotinylation*. Female *Pitx3*^*IRES2-*^ *tTA/+* mouse VTA were bilaterally injected with either AAV9-TRE-eGFP or -shRit2. Brains were harvested 4-5 weeks post-injection, and DAT surface expression was measured in *ex vivo* striatal slices by surface biotinylation as described in *Materials and Methods*, following treatment ±1μM PMA, 30 min, 37°C. Representative blots are shown in the top of each panel, and average data are presented at the bottom of each panel. DAT surface levels are expressed as %total DAT, ±S.E.M, n=5-8 slices from n=3 independent mice/virus. **A.** *Dorsal striatum*. Neither PKC activation nor Rit2-KD had an effect on DAT surface expression (two-way ANOVA: Interaction: F_(1, 22)_ = 0.051, p = 0.82; drug: F_(1, 22)_ = 0.58, p = 0.46; virus: F_(1, 22)_ = 0.18, p = 0.68). **B.** *Ventral striatum.* PKC activation and Rit2-KD significantly decreased DAT surface expression, and Rit2-KD blocked PKC-stimulated DAT internalization (two-way ANOVA: Interaction: F_(1, 18)_ = 4.54, p = 0.047; drug: F_(1, 18)_ = 4.25, p = 0.054; virus: F_(1, 18)_ = 7.28, p = 0.015. Sidak’s multiple comparisons test: eGFP(veh) vs eGFP(PMA): *p = 0.04; eGFP(veh) vs shRit2(veh): *p = 0.02; shRit2(veh) vs shRit2(PMA): p > 0.99).

Our previous study probed the impact of DAergic Rit2 KD on basal DAT surface levels in males (39). Therefore, we next asked whether Rit2 was required for PKC-stimulated DAT internalization in male DS and VS. Similar to females, PKC activation had no effect on DAT surface expression in DS, but significantly decreased DAT surface expression in VS, measured in control (eGFP-injected) male mice (Fig 4A). Also similar to females. DAergic Rit2-KD completely abolished PKC-stimulated DAT internalization in male VS (Fig. 4B). Given that phorbol esters can stimulate a variety of signaling pathways in addition to PKC, we further tested whether PMA-induced DAT internalization in VS was PKC-mediated. Pre-treatment with the PKC-specific inhibitor BIM I (1µM, 15 min, 37°C) significantly abolished PMA-induced DAT internalization, and treatment with BIM I alone was not significantly different from BIM I/PMA (Fig. 4C), clearly demonstrating that PMA-mediated DAT internalization in DAergic terminals is PKC-dependent.

**Figure 4.**
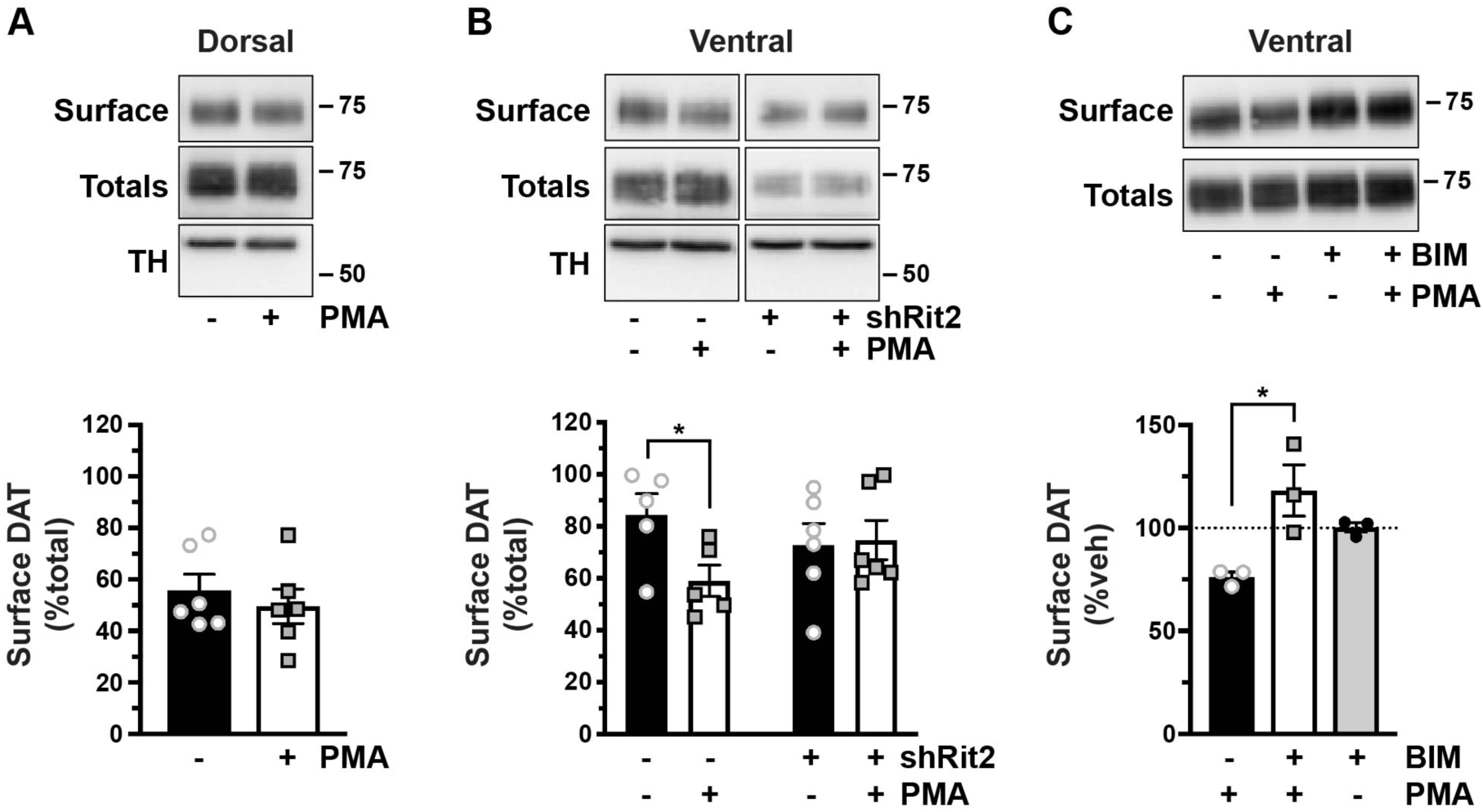
PKC-induced DAT internalization in males is limited to ventral striatum and requires Rit2. *Conditional Rit2 silencing in DA neurons and ex vivo striatal slice biotinylation*. Male *Pitx3*^*IRES2-tTA/+*^ mouse VTA were bilaterally injected with either AAV9-TRE-eGFP or -shRit2. Brains were harvested 4-5 weeks post-injection, and DAT surface expression was measured in *ex vivo* striatal slices by surface biotinylation as described in *Materials and Methods*, following treatment ±1μM PMA, 30 min, 37°C. Representative blots are shown in the top of each panel, and average data are presented at the bottom of each panel. **A.** ***Effect of PKC activation on DAT surface levels in dorsal vs. ventral** striatum*. DAT surface levels were measured in AAV9-TRE-eGFP-injected mice and are expressed as %total DAT ±S.E.M. PKC activation had no effect on DAT surface expression in dorsal striatum, but significantly decreased DAT surface levels in ventral striatum (two-way ANOVA: Interaction: F_(1, 18)_ = 1.96, p = 0.18; region: F_(1, 18)_ = 7.76, p = 0.01; drug: F_(1, 18)_ = 5.30, p = 0.03. Sidak’s multiple comparisons test (vehicle vs. PMA): *Dorsal:* p = 0.76; *Ventral:* p = 0.043 n = 5-6 slices from 3 independent mice. **B.** *Ventral striatum*. DAT surface levels are expressed as %total DAT ±S.E.M. PKC activation significantly decreased DAT surface expression in male VS (*p = 0.037, two-tailed Student’s t test, n = 5 slices from 3 independent mice per condition). Rit2-KD significantly blocked PKC-stimulated DAT internalization in ventral striatum (shRit2: p = 0.86, two-tailed Student’s t test, n=6 slices from 3 independent mice). **C.** *PMA-induced DAT internalization is PKC-mediated*. DAT surface expression was measured in *ex vivo* VS slices prepared from C57Bl/6J mice and were pretreated ±1µM BIM I, 15 min, 37°C prior to PMA treatment as described above. DAT surface levels are expressed as %vehicle ±S.E.M. BIM I pre-treatment significantly abolished PMA-mediated DAT surface loss (one-way ANOVA F_(2, 6)_ = 8.08, p = 0.02; Sidak’s multiple comparisons test, PMA vs BIM/PMA: *p = 0.01, BIM/PMA vs BIM: p = 0.25, n = 3 independent mice per condition).

### Releasing the PKC-sensitive DAT endocytic brake drives DAT-Rit2 dissociation at the plasma membrane

We next sought to decipher the molecular mechanisms by which Rit2 impacts DAT trafficking. We first asked whether driving DAT internalization, by disengaging the PKC-sensitive DAT endocytic brake, impacts the DAT-Rit2 interaction at the plasma membrane. To specifically interrogate the DAT surface population and its associated protein complex, we leveraged a BBS-targeted surface labeling strategy (46-48) to label DAT in intact cells. We engineered a BBS into the DAT extracellular loop 2, a site we previously successfully targeted for bio-orthogonal DAT labeling (49), and which also tolerates an HA epitope (50). BBS-DAT expressed and functioned comparably to wt-DAT, and PKC activation acutely decreased BBS-DAT function to 67.97 ±5.9% control levels (Fig. 5A), which is comparable to PKC-mediated DAT downregulation as previously reported by our group and others (22,38,43,49,51). We first tested whether BBS-DAT could: 1) specifically isolate surface DAT via bungarotoxin labeling and pulldown, and 2) recover DAT-associated proteins. HEK293T cells expressing HA-Rit2 and either BBS-DAT or wt-DAT were incubated with α-BTX-b, and DAT surface complexes were isolated from cell lysates by streptavidin pulldown. We specifically recovered BBS-DAT, but not wt-DAT, following α-BTX-b incubation (Fig. 5B), demonstrating the selectivity of the BBS pulldown approach to label and isolate surface BBS-DAT. Importantly, Rit2 was recovered from pulldowns with BBS-DAT, but not in control pulldowns from cells expressing wt-DAT, demonstrating that Rit2 is recovered specifically following surface DAT pulldown. Additionally, BBS-DAT pulldowns did not recover the Rit2 homolog, Rit1 (Fig. 5C), consistent with a specific association between DAT and Rit2. We further asked whether other proteins required for PKC-mediated brake release are also part of the DAT surface complex. We previously reported that the non-receptor tyrosine kinase, Ack1 (TNK2) imposes the PKC-sensitive endocytic brake, and that PKC-mediated Ack1 inactivation is required for PKC-stimulated DAT internalization (24). However, it is not known whether Ack1 is associated with DAT at the plasma membrane. Following α-BTX-b labeling and pulldown, Ack1 was recovered from cells expressing BBS-DAT, but not from control cells expressing wt-DAT (Fig. 5D), demonstrating that Ack1 is part of the DAT surface complex. Thus, BBS-DAT has precise utility to interrogate surface DAT and its associated proteins, such as Rit2 and Ack1.

**Figure 5.**
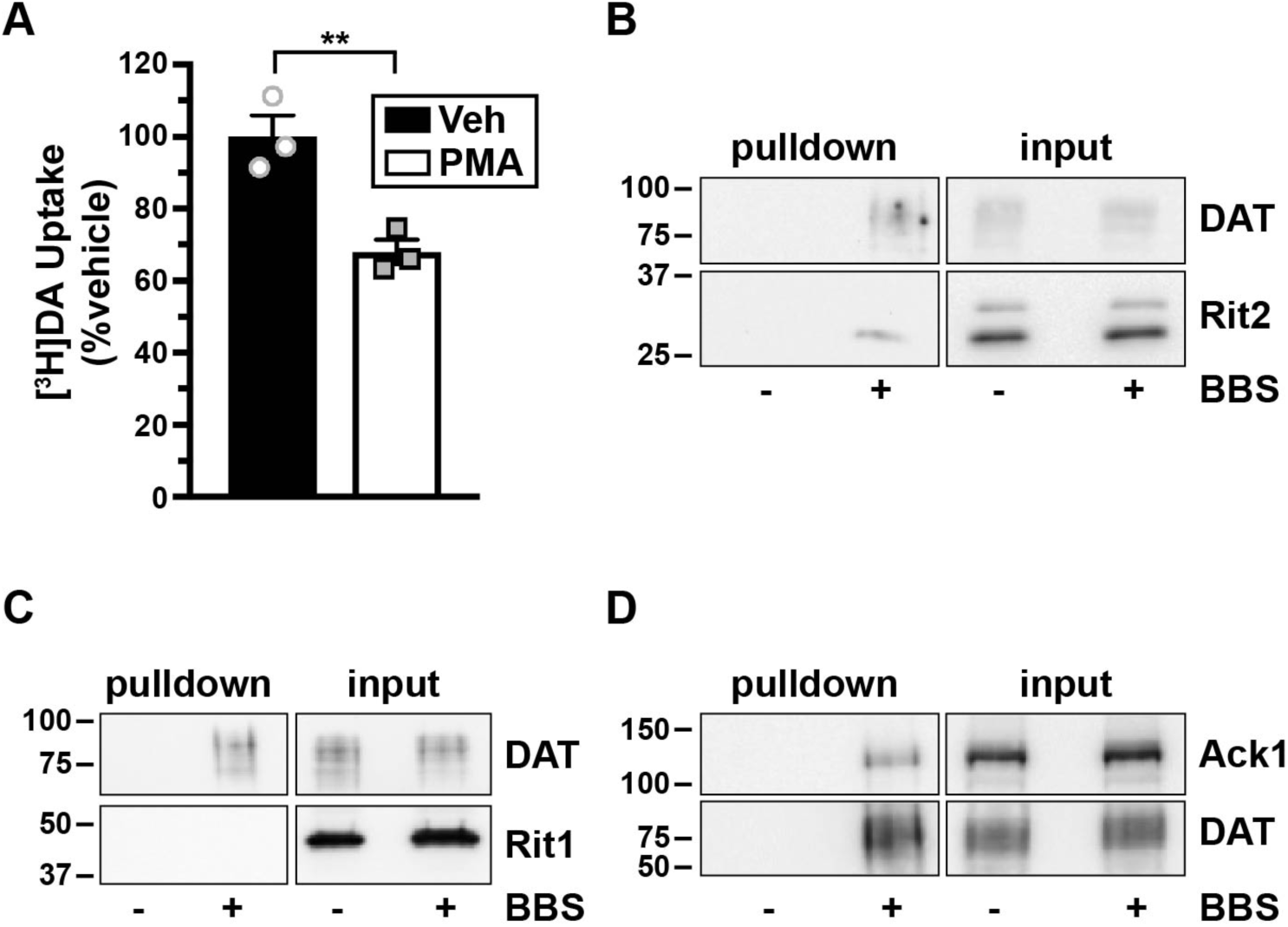
Surface DAT associates with Rit2 and Ack1, but not Rit1. **A.** [^*3*^*H]DA uptake assay.* SK-N-MC cells expressing BBS-DAT were treated ±1µM PMA, 30 min, 37°C and [^3^H]DA uptake was measured as described in *Materials and Methods*. Average data are expressed as %vehicle-treated specific [^3^H]DA uptake ±S.E.M. **p=0.009, two-tailed Student’s t test, n=3. **B-D**. *BBS-DAT pulldowns*. HEK293T cells were co-transfected with DAT (±BBS tag), and either HA-Rit2 **(B)**, GFP-Rit1 **(C)**, or Ack1-HA **(D)** and DAT surface complexes were labeled and isolated by streptavidin pulldown as described in *Materials and Methods.* Representative immunoblots for pull-downs (left panels) and their respective inputs (1/4 total input, right panels) are presented (n=3 independent experiments).

Given our previous cellular imaging results (38), we hypothesized that PKC activation causes DAT and Rit2 to dissociate. Since PKC-stimulated DAT internalization can only occur when the endocytic brake is disengaged, we first leveraged BBS-DAT pulldowns to ask whether PKC-mediated brake release alters the DAT-Rit2 surface association. PKC activation (1µM PMA, 30 min, 37°C) significantly decreased the DAT-Rit2 plasma membrane association (Fig. 6A). Additionally, DAT and Rit2 significantly dissociated when we directly released the DAT endocytic brake, by inactivating Ack1 with AIM-100 (20µM, 30 min, 37°C) (Fig. 6B). DAT and Rit2 may specifically dissociate at the cell surface in response to releasing the PKC-sensitive endocytic brake, or may do so following any stimulus that drives DAT internalization. To discern between these two possibilities, we measured the DAT-Rit2 surface association in response to AMPH treatment, which also accelerates DAT internalization, but is Rho-dependent (52). In contrast to PKC-stimulated DAT-Rit2 dissociation, AMPH treatment (10µM, 30 min, 37°C) significantly increased the DAT-Rit2 surface association (Fig. 6C). Thus, the DAT-Rit2 surface dissociation occurs specifically when the PKC-sensitive DAT endocytic brake is disengaged, either in response to PKC activation or direct Ack1 inactivation, but is not a general result of accelerated DAT endocytosis.

**Figure 6.**
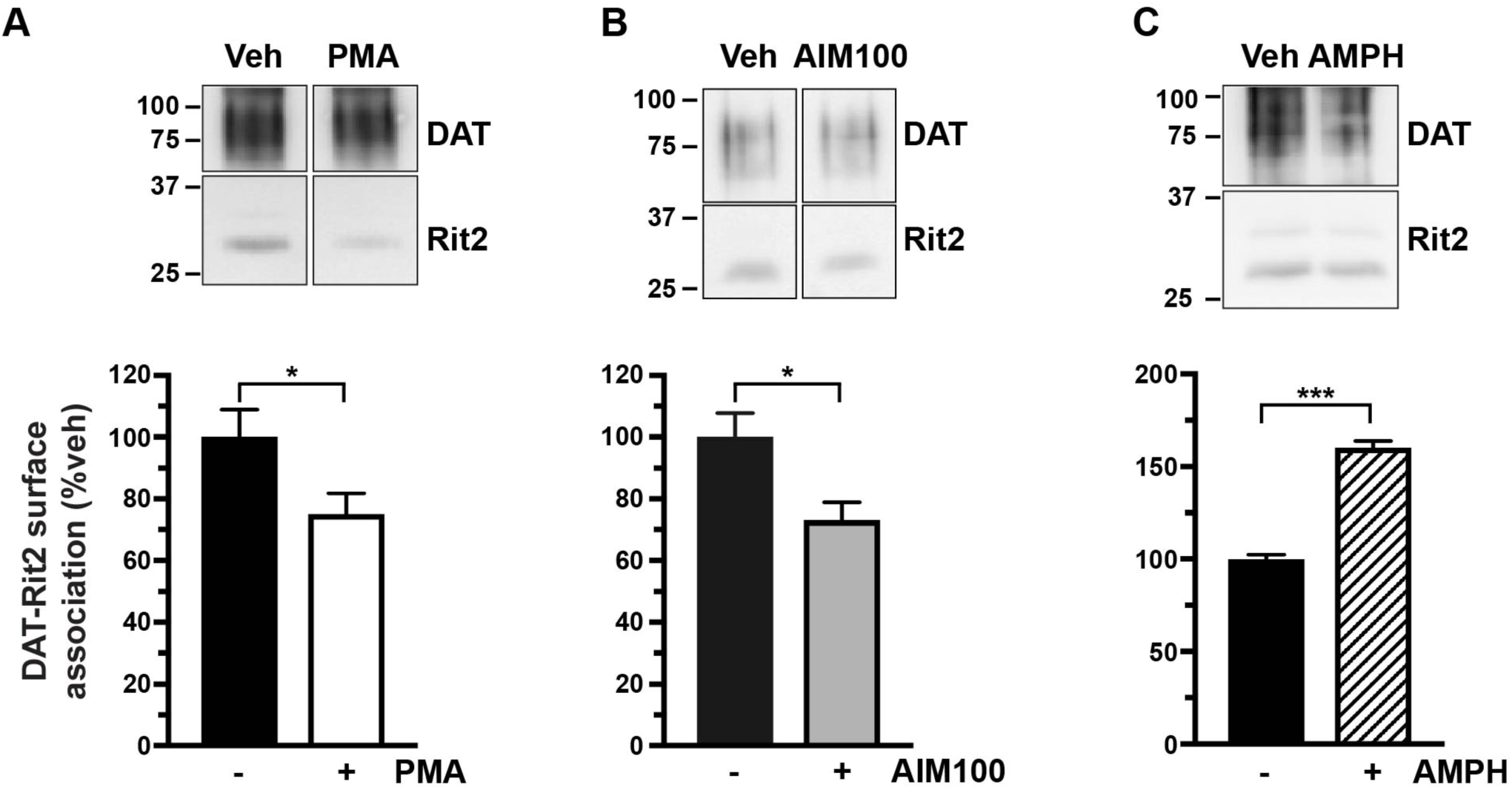
PKC-mediated endocytic brake release drives DAT-Rit2 dissociation at the plasma membrane. *BBS-DAT pull-downs*. HEK293T cells were co-transfected with BBS-DAT and HA-Rit2, were treated ±the indicated drugs, 30 min, 37°C, labeled with α-BTX-b, and DAT surface complexes were isolated as described in *Materials and Methods. Top panels*: Representative immunoblots. *Bottom panels:* Average DAT-Rit2 association expressed as %vehicle-treated control ±S.E.M., assessed by two-tailed Student’s t test. **A.** PKC activation significantly decreased the DAT-Rit2 surface association, *p=0.04, n=6. **B**. Ack1 inactivation with AIM-100 decreased the DAT-Rit2 surface association, *p=0.03, n=4. **C**. AMPH treatment significantly increased DAT-Rit2 surface association, ***p=0.0002, n=3.

### The DAT amino terminus is integral to the DAT-Rit2 interaction and PKC-stimulated dissociation

The DAT-Rit2 interaction was originally identified in a yeast 2-hybrid screen, using DAT C-terminal residues 587-596 (FREKLAYAIA) as bait (38). However, it is not known which DAT domains are required (or sufficient) for the DAT-Rit2 association in the context of intact DAT protein, nor is it known whether any DAT domains are specifically required for PKC-stimulated DAT-Rit2 dissociation. Rit2 specifically binds DAT, but not SERT (38), thus we hypothesized that replacing DAT N- or C-termini with those of SERT may define DAT domains required for Rit2 binding and/or PKC-stimulated DAT-Rit2 dissociation. To test this possibility, we leveraged a series of DAT/SERT chimeras we previously characterized (53), in which either the DAT N-, C-, or both termini were substituted with those of SERT. HEK293T cells were co-transfected with YFP-Rit2 and CFP-tagged versions of these chimeras, and we performed live FRET imaging to quantify the chimera-Rit2 interactions. As we previously reported, control CFP-DAT and YFP-Rit2 elicited a significant FRET signal as compared to soluble YFP/CFP expression (Fig. 7A). Interestingly, replacing the DAT N-terminus with that of SERT (CFP-N-S/DAT) significantly increased the DAT-Rit2 interaction compared to CFP-DAT, whereas replacing the DAT C-terminus (CFP-DAT/C-S) or both termini (CFP-S/DAT/S) did not affect the DAT-Rit2 interaction (Fig. 7A). We also observed a significant increase in the interaction between YFP-Rit2 and CFP-N-S/DAT using the donor recovery after photobleaching approach (Fig. 7B), demonstrating that the FRET signal is a bona fide interaction between the fluorophores. Using the BBS pulldown approach, Rit2 was likewise recovered with BBS-tagged versions of each DAT/SERT chimera (Fig. 7C). We next asked whether PKC-stimulated DAT-Rit2 dissociation requires either the DAT N- and/or C-termini. Substituting the DAT C-terminus with that of SERT (DAT/C-S) had no significant effect on PKC-stimulated DAT-Rit2 dissociation, as compared to wt-DAT controls (Fig. 7D, one-way ANOVA with Dunnett’s multiple comparison test, p=0.69). However, substituting the DAT N-terminus with that of SERT (N-S/DAT) completely abolished PKC-stimulated DAT/Rit2 dissociation, and there was a strong trend for attenuated DAT/Rit2 dissociation when both DAT N- and C-termini were replaced by SERT (S/DAT/S) (Fig 7D, p=0.058). Taken together, these results indicate that the DAT N-terminus is required for the PKC-stimulated DAT-Rit2 dissociation, and that the SERT N-terminus does not suffice. However, SERT N- and C-termini suffice to maintain the DAT-Rit2 association.

**Figure 7.**
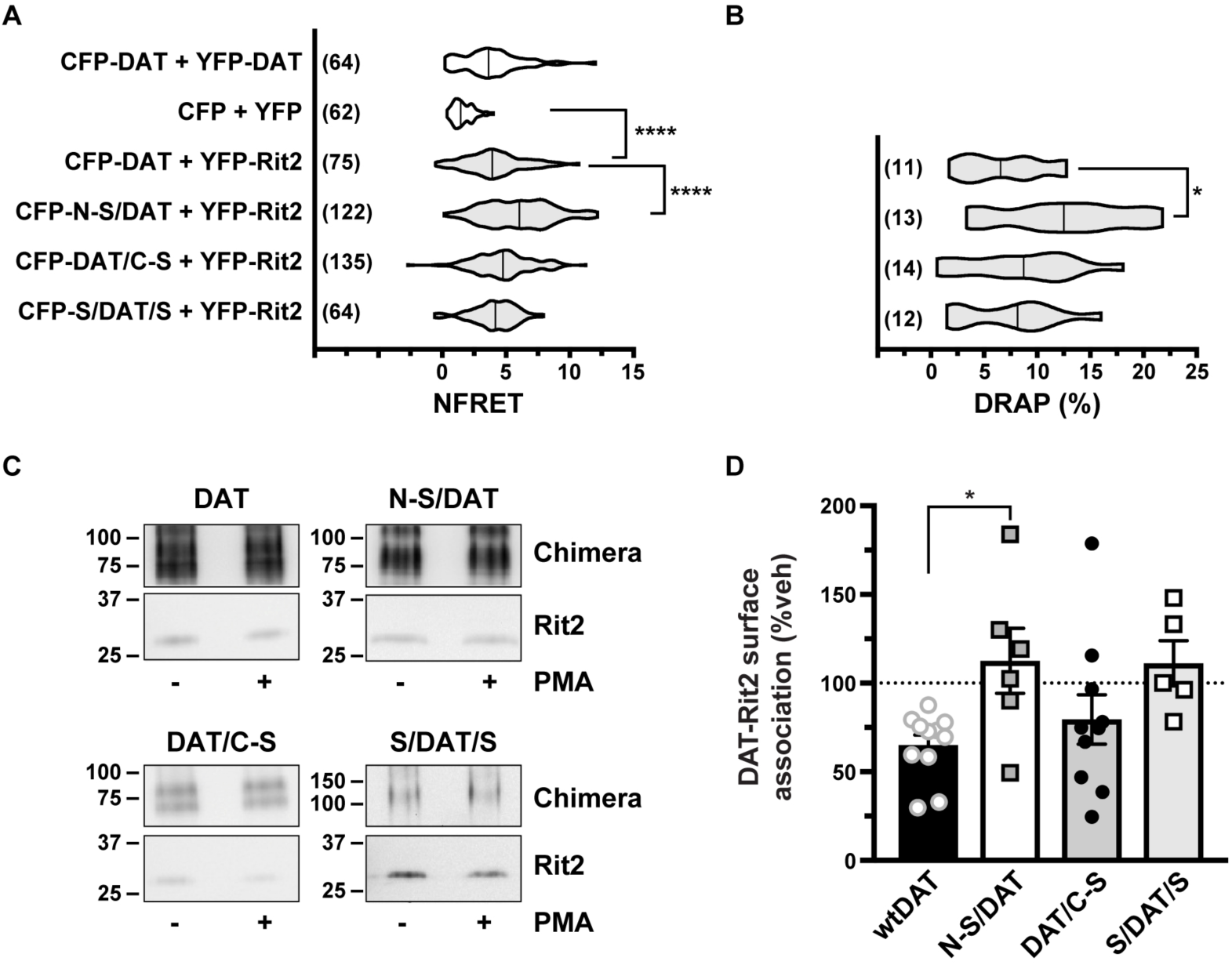
The SERT N-terminus promotes the DAT-Rit2 interaction and blocks PKC-stimulated DAT-Rit2 dissociation. **A, B.** *FRET studies.* HEK293 cells were transfected with the indicated plasmids and surface FRET measurements were made 24 hrs. post-transfection, as described in *Materials and Methods*. Average NFRET values (x100) for the indicated pairs are presented as violin plots, with median lines provided within each violin. N values are provided for each violin. **A.** NFRET values. Asterisks indicate significant differences between the indicated pairs (Kruskal-Wallis test, p<0.0001 with Dunn’s multiple comparisons test: DAT vs N-S/DAT: ****p < 0.0001; vs DAT/C-S: p = 0.13; vs S/DAT/S: p > 0.99; vs CFP/YFP: ****p < 0.0001. **B.** *DRAP values*. one-way ANOVA F_(3, 46)_ = 3.31, p = 0.028; Dunnett’s multiple comparisons test: DAT vs N-S/DAT: *p = 0.02; DAT/C-S: p = 0.81; vs S/DAT/S: p = 0.99.) **C, D.** *BBS-DAT pull-downs.* HEK293T cells were co-transfected with HA-Rit2 and the indicated BBS-tagged proteins. Cells were treated ±1μM PMA, 30 min, 37°C, surface-labeled with α-BTX-b and DAT surface complexes were recovered by streptavidin pulldown as described in *Materials and Methods*. **C.** Representative immunoblots. **D**. Average data presented as %vehicle-treated DAT-Rit2 association for each indicated protein. N-S/DAT significantly blocked PKC-stimulated DAT-Rit2 dissociation (one-way ANOVA F_(3, 28)_ = 3.44, p = 0.03; Dunnett’s multiple comparisons test: DAT vs N-S/DAT: *p = 0.03; vs DAT/C-S: p = 0.69; vs S/DAT/S: p = 0.06, n=5-11).

### The DAT amino terminus is required for PKC-stimulated DAT internalization

Since the DAT N-terminus is required for PKC-stimulated DAT-Rit2 dissociation, this raised the possibility that the N-terminus may also be required for stimulated DAT internalization, driven by release of the DAT endocytic brake. To test these possibilities, we measured DAT and DAT/SERT chimera internalization rates in response to either PKC activation or direct Ack1 inactivation (with AIM-100) in stably-transfected SK-N-MC cells, in which we previously characterized both PKC- and AIM-100-stimulated DAT internalization (24,26). PKC activation (1µM PMA, 10 min, 37°C) significantly increased wt-DAT internalization, and substituting the DAT C-terminus with the SERT C-terminus (DAT/C-S) did not significantly affect PKC-stimulated internalization (Fig. 8A). However, PKC-stimulated DAT internalization was abolished when either the DAT N-terminus, or both N- and C-termini, were substituted with SERT termini (N-S/DAT and S/DAT/S, Fig. 8A). In contrast, direct Ack1 inactivation (20µM AIM-100, 10 min, 37°C) significantly stimulated wt-DAT, N-S/DAT, and DAT/C-S internalization, but had no effect on S/DAT/S internalization (Fig. 8A). We further evaluated the basal endocytic rates of the chimeric DATs, as compared to wt-DAT. As presented in Figure 8B, none of the chimera basal internalization rates differed significantly from wt-DAT. To assure that these effects were not due to the lack of Rit2 expression in SK-N-MC cells, we further assessed basal and PKC-stimulated N-S/DAT internalization in stably transfected SK-N-DZ cells, which endogenously express Rit2 (Table I). In SK-N-DZ cells, N-S/DAT internalized significantly slower than wt-DAT under basal conditions (Fig 8C). Additionally, while PKC activation significantly increased the wt-DAT internalization rate, it had no effect on N-S/DAT internalization as compared to its own vehicle control (p=0.17, one-way ANOVA with Bonferroni’s multiple comparison test, n=4-7). These results demonstrate that the DAT N-terminus is required, and that the SERT N-terminus does not suffice, for PKC-stimulated DAT internalization.

**Figure 8.**
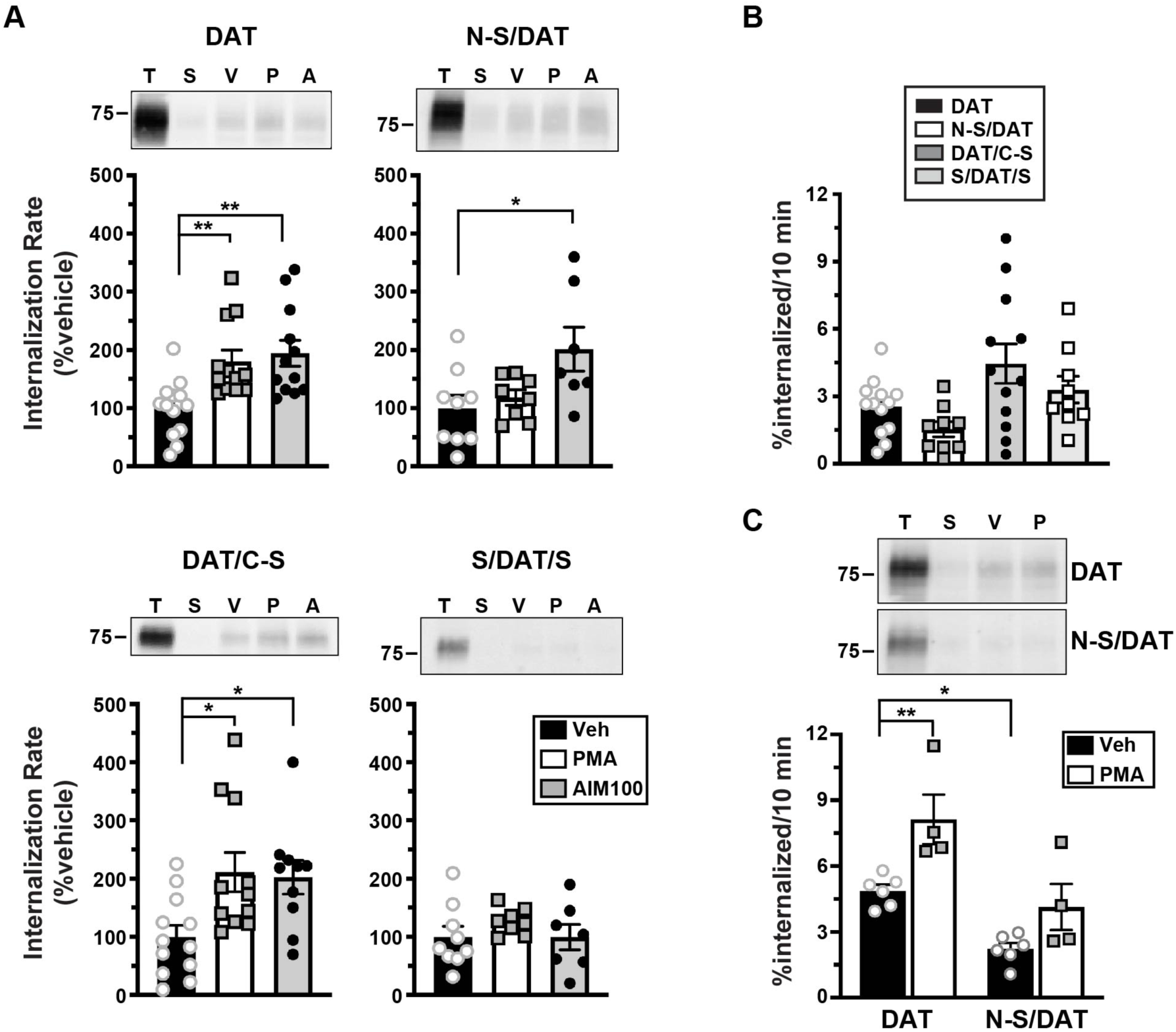
The DAT N-terminus is required for PKC-stimulated internalization. *DAT Internalization assays*. Wt-DAT and DAT/SERT chimera internalization rates were measured ±1μM PMA or ±20µM AIM-100, 10 min, 37°C, as described in *Materials and Methods. Tops:* Representative immunoblots showing total surface expression at t=0 (T), strip control (S), and internalized protein during either vehicle (V), PMA (P), or AIM-100 (A) treatments. *Bottoms:* Averaged data. **A**. Stimulated internalization in stable SK-N-MC cell lines. Rates are expressed as %vehicle-treated controls ±S.E.M. Asterisks indicate a significant difference from vehicle controls (one-way ANOVA with Dunnett’s multiple comparisons test, n=8-13. **<u>DAT</u>**: ANOVA F_(2, 34)_ = 7.94, p = 0.0015; veh vs PMA: **p = 0.007; veh vs AIM-100: **p = 0.002. **<u>N-S/DAT</u>**: ANOVA F_(2, 21)_ = 4.38, p = 0.03; veh vs PMA: p = 0.82; veh vs AIM-100: *p = 0.02. **<u>DAT/C-S</u>**: ANOVA F_(2, 30)_ = 5.22, p = 0.01; veh vs PMA: *p = 0.01; veh vs AIM-100: **p = 0.026. **<u>S/DAT/S</u>**: ANOVA F_(2, 21)_ = 0.84, p = 0.44. **B.** *Basal internalization in stable SK-N-MC cell lines*. Chimera basal internalization rates did not significantly differ from wt-DAT (one-way ANOVA F_(3, 39)_ = 4.046, p = 0.013; Dunnett’s multiple comparisons test: DAT vs: N-S/DAT: p = 0.54; vs DAT/C-S: p = 0.06; vs S/DAT/S: p = 0.72, n=9-12). **C.** *DAT and N/S-DAT internalization in stable SK-N-DZ cell lines*. Average internalization rates are expressed as %surface protein internalized/10 min, ±S.E.M. Asterisks indicate a significant difference from the indicated protein or treatment (two-way ANOVA: interaction: F_(1, 16)_ = 1.06, p = 0.32; construct: F_(1, 16)_ = 15.6, p = 0.001; drug: F_(1, 16)_ = 25.6, p = 0.0001; Tukey’s multiple comparisons test: DAT(veh) vs DAT(PMA): *p = 0.01, DAT(veh) vs N-S/DAT(veh): *p = 0.03; N-S/DAT(veh) vs NS/DAT(PMA): p = 0.21, n=4-7).

### Rit2 and Ack1 independently converge on DAT in response to PKC

We next asked whether there is a mechanistic relationship between Rit2, Ack1, and PKC-mediated DAT endocytic brake release. Rit2 may be either upstream or downstream from Ack1 in the signaling cascade that leads from PKC to DAT. Alternatively, Rit2 and Ack1 may respond independently to PKC activation to stimulate DAT internalization. We first used Rit2-KD in SK-N-DZ cells to ask whether Rit2 was required for PKC-mediated Ack1 inactivation, a requisite step for PKC-mediated release of the endocytic brake and for stimulated DAT internalization (24). As we previously reported, in vector-transduced cells, PKC activation significantly reduced pY284-Ack1, to levels comparable to that achieved with the Ack1 inhibitor, AIM-100 (Fig. 9A). In cells transduced with shRit2-107, pY284-pAck1 levels, were also significantly reduced in response to either PKC activation or AIM-100 treatment, and there was no difference in the magnitude change of pY284-pAck1 levels following PKC activation (Fig. 9A). These results indicate that Rit2 is not likely upstream of Ack1 in the signaling pathway that leads from PKC to Ack1 inactivation.

**Figure 9.**
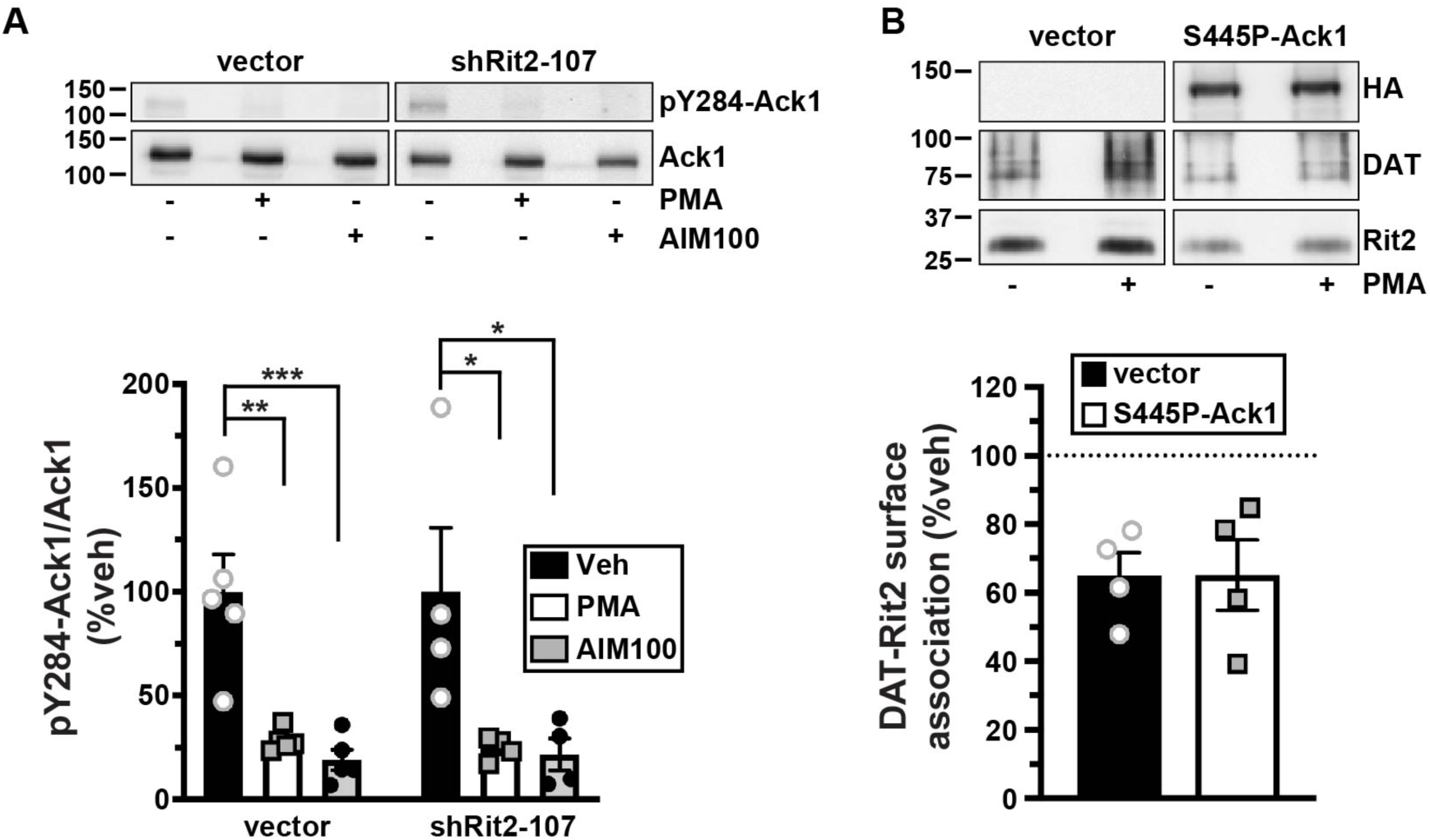
Rit2 and Ack1 independently converge on DAT downstream of PKC activation. **A.** *Effect of Rit2 silencing on PKC-mediated Ack1 inactivation*. SK-N-DZ cells were transduced with lentiviral particles expressing either vector (pGIPZ) or shRit2-107, and were treated ±1μM PMA or ±20μM AIM-100, 10 min, 37°C. pY284Ack1 and total Ack1 levels were measured from parallel lysate aliquots by immunoblot. *Top*: Representative immunoblots. *Bottom*: Average pY284-Ack1 levels expressed as %vehicle-treated control cells ±S.E.M. Asterisks indicate a significant difference from vehicle-treated controls (one-way ANOVA with Dunnett’s multiple comparisons test, **<u>vector:</u>** ANOVA F_(2, 12)_ = 16.43, p = 0.0004; veh vs PMA: **p= 0.001; vs AIM-100: ***p = 0.0004; **<u>shRit2-107</u>**: ANOVA F_(2, 9)_ = 5.858, p = 0.02; veh vs PMA: *p =0.03; vs AIM-100: *p = 0.03, n=4-5). **B**. *PKC-stimulated DAT-Rit2 dissociation*. HEK293T cells were triple-transfected with BBS-DAT, HA-Rit2, and either empty vector or S445P-Ack1-HA. Cells were treated±1μM PMA, 30 min, 37°C, labeled with α-BTX-b, and DAT complexes were recovered by streptavidin pulldown as described in *Materials and Methods. Top*: Representative immunoblots. *Bottom*: Average DAT-Rit2 association expressed as %vehicle-treated control ±S.E.M. p=0.99, two-tailed Student’s t test, n=4.

We next asked whether PKC-mediated Ack1 inactivation is required for, and therefore upstream of, PKC-stimulated DAT-Rit2 dissociation. To test this possibility, we measured DAT-Rit2 dissociation in cells co-transfected with DAT, Rit2, and either vector or the PKC-insensitive, constitutively active Ack1 mutant (S445P) which we previously reported blocks PKC-stimulated DAT internalization (24). In control cells, PKC activation drove a significant DAT-Rit2 dissociation (Fig. 9B). In cells co-transfected with S445P-Ack1, PKC activation likewise drove DAT-Rit2 dissociation, at levels indistinguishable from vector controls (Fig. 9B). These results demonstrate that PKC-stimulated DAT-Rit2 dissociation does not require Ack1 inactivation. Moreover, they demonstrate that even in conditions where DAT cannot internalize in response to PKC activation (i.e. because Ack1 is constitutively active), PKC activation still drives DAT and Rit2 to dissociate. Thus, PKC-stimulated DAT-Rit2 dissociation is likely to occur prior to Ack1-mediated release of the DAT endocytic brake.

## DISCUSSION

DAT is stabilized at the cell surface by a PKC-sensitive endocytic brake that requires 1) residues in both the DAT N- and C-termini to engage (23,27), and 2) Ack1 inactivation to be released (24). Here, we found that Rit2 is required to release the PKC-sensitive DAT endocytic brake in cell lines and tissues where Rit2 is endogenously expressed, such as SK-N-DZ cells (Fig. 2C) and in VS (Fig. 3 and Fig. 4), respectively. Curiously, over two decades of evidence has demonstrated that PKC activation stimulates DAT internalization in in a wide range of cultured cell lines (42,50,54-56), which our results demonstrate have negligible, if any, Rit2 expression (Table 1). Given that Ack1 is ubiquitously expressed, and that Rit2 and Ack1 converge independently on DAT in response to PKC activation (Fig. 9), we conclude that PKC-stimulated Ack1 inactivation can suffice to release the endocytic brake when DAT is heterologously expressed in a context that does not express Rit2. However, when DAT is expressed in its appropriate context (i.e. striatal terminals), Rit2 expression is absolutely required for PKC-stimulated DAT internalization, and PKC-mediated Ack1 inactivation alone does not suffice to release the endocytic brake (Fig. 3 and Fig. 4). These results further suggest that although Rit2 and Ack1 independently converge on DAT in response to PKC activation in cell lines (Fig. 9), there may be an, as of yet, unidentified DAergic-specific mechanism(s) that facilitate a Rit2/Ack1 interdependence required for endocytic brake release.

**Table I.** Rit2 expression in mammalian cell lines and rodent brain regions. Rit2 mRNA expression was determined by RT-qPCR and normalized to internal GAPDH values. n = 2-4

| Species/Cell line | *Rit2* ($2^{-\Delta C_t} \times 10^4 \pm \text{S.E.M}$) |
| --- | --- |
| **Mouse** |  |
| CAD | 0.02 ± 0.0067 |
| N2a | 0.09 ± 0.052 |
| NIH/3T3 | No signal detected |
| Midbrain | 72.0 ± 12.8 |
| Cortex | 54.8 ± 9.19 |
| **Rat** |  |
| AN27 | No signal detected |
| PC12 | No signal detected |
| Midbrain | 110.0 ± 0.51 |
| **Human** |  |
| HEK293T | 0.40 ± 0.34 |
| SH-SY5Y | 1.3 ± 0.56 |
| SK-N-MC | 0.09 ± 0.07 |
| SK-N-DZ | 27.2 ± 18.7 |

Our previous study using total (i.e. not sub-dissected) striatal slices found that PKC activation decreased DAT surface expression by ∼20% (26). Interestingly, in the current study, PKC activation drove ∼35% DAT surface loss in both male and female VS, but had no effect on DAT surface expression in DS from either sex (Fig. 3). These data suggest that the somewhat modest PKC-stimulated DAT surface loss detected in total striatum reflects robust DAT surface loss in VS, diluted by the lack of a net effect in DS. Is DAT endocytosis completely resistant to PKC activation in DS? It is currently unknown which PKC isoform(s) stimulate DAT internalization. We activated PKC with the phorbol ester PMA, which activates all diacylglycerol-sensitive PKC isoforms, including PKC-α, β1, β2, γ, and δ. PKCβ activity is required for D2 receptor-mediated DAT insertion into the plasma membrane, and selective PKCβ activation rapidly delivers DAT to the cell surface in both cell lines and striatal synaptosomes (57). Thus, it is possible that in DS, Rit2-dependent, PKC-stimulated DAT internalization may be countered by PKCβ-mediated DAT insertion, resulting in no detectable net change within the 30-min timeframe we tested. In contrast to DS, DAT surface levels in VS were significantly decreased in response to PMA treatment, suggesting that the balance between PKCβ-mediated DAT insertion and Rit2-dependent DAT internalization may differ between DS and VS. A recent study by Blakely and colleagues (58) further supports this premise, in which they reported that D2R activation increased DAT surface expression in the DS, but not VS. Given that few studies, to date, have identified the receptor-mediated signaling pathways that lead to PKC-stimulated DAT internalization, nor the temporal profile of DAT response to PKC activation in DAergic terminals, it remains unclear how DAT insertion and internalization balance may occur. Finally, although our current data indicate that Rit2-KD is required for PKC-stimulated DAT endocytosis in female and male VS, it is important to bear in mind that DAT basal surface expression in both male (39) and female (Fig. 3B) VS were reduced following DAergic Rit2-KD. Thus, there is a possibility that a general floor effect blocked PKC-mediated DAT internalization, rather than a requirement for Rit2. However, we do not believe that this is likely, as DAT is capable of undergoing additive degrees of internalization when cells are subjected to dual PKC activation and AMPH exposure (59).

We used an extracellular BBS-tag to interrogate surface DAT and its associated protein complex. This approach was previously used successfully by several groups (46,47,60), and has distinct advantages over co-IP approaches, since 1) the α-BTX/BBS affinity is significantly higher than that of antibody/antigen interactions, and 2) α-BTX/BBS binding is maintained in detergent lysates. Moreover, using an extracellular labeling approach, in general, maintains intracellular protein complexes that might be disrupted using an intracellularly-targeted antibody in lysates/solution. Interestingly, although we observed significant DAT-Rit2 dissociation in response to releasing the PKC-sensitive DAT endocytic brake (Fig. 6), AMPH-stimulated DAT internalization increased the surface DAT-Rit2 population. Surface DAT is distributed among several membrane microdomains (38,61-67). Moreover, we previously reported that, 1) PKC activation preferentially depletes DAT from cholera toxin-positive (Ctx+) microdomains (26), and 2) there is significantly more DAT-Rit2 co-localization in Ctx+ microdomains (38). Conversely, AMPH treatment increases DAT localization to Ctx+ domains (63). Taken together, these results suggest that DAT may dissociate from Rit2, and internalize, preferentially from Ctx+ microdomains in response to PKC activation. In contrast, AMPH potentially drives DAT internalization from Ctx-domains, in which there is less DAT-Rit2 interaction, thereby leaving an enriched DAT-Rit2 population at the cell surface. We originally identified the DAT-Rit2 interaction via a yeast 2-hybrid screen using the C-terminal DAT domain “FREKLAYAIA” as bait, and FRET studies revealed that Rit2 directly interacts with DAT, but not SERT, at the plasma membrane (38). However, the domains required for the DAT-Rit2 interaction, and their requirement for either PKC-stimulated DAT internalization or DAT-Rit2 dissociation were not defined. To address these questions, we used a series of DAT/SERT chimeras previously reported by our lab (53), and found that the DAT N-terminus was required for PKC-stimulated DAT-Rit2 dissociation (Fig. 8). Indeed, under basal conditions, N-S/DAT and Rit2 interacted to a significantly higher degree than DAT-Rit2 controls (Fig. 6), consistent with a lack of DAT-Rit2 dissociation for the N-S/DAT chimera and higher steady-state interaction. N-S/DAT also basally internalized significantly slower than wt-DAT selectively in SK-N-DZ, consistent with its inability to disengage the endocytic brake. Moreover, DAT/C-S retained both Rit2 interaction and ability release the endocytic brake in response to PKC activation. This was surprising, given that 1) the DAT C-terminal bait (FREKLAYAIA) used to identify the DAT/Rit2 interaction is highly conserved across the SLC6 gene family (23,44), and 2) full-length SERT does not interact with Rit2 (38). So, while the FREKLAYAIA domain is sufficient to interact with Rit2, its context within full-length DAT or SERT appears to dictate ultimate Rit2 binding potential. Interestingly, although N-S/DAT was resistant to PKC-stimulated internalization, it retained AIM-100-dependent internalization, whereas S/DAT/S did not (Fig. 8), indicating that the DAT N-terminus is not required for direct Ack1-dependent endocytic brake release, and further supports the hypothesis that Rit2 and Ack1 converge on DAT independently in response to PKC activation. It should also be noted that we, and others, reported that AIM-100 also binds non-competitively to DAT (24,68), and recent reports suggest that AIM-100 can enhance DAT surface oligomerization (68,69). Since both Ack1-dependent and AIM-100-stimulated internalization are specific for DAT, and not SERT, it is possible that S/DAT/S endocytic resistance to AIM-100 may be because the substituted SERT domains perturb the DAT/AIM-100 interaction. Likewise, it is possible that the ability of N-S/DAT to internalize in response to AIM-100, but not PKC activation, may be due to a direct AIM-100 effect on DAT.

In this study we present one of the first descriptions of region- and sex-dependent differences in DAT trafficking regulation. Furthermore, we greatly extend our knowledge of the mechanisms by which Rit2 governs DAT surface expression in bona fide DAergic terminals. Future studies that examine the cell autonomous endogenous signaling events that drive striatal DAT trafficking, and require Rit2, will shed further light on mechanisms that influence DA clearance and DA-dependent behaviors.

## EXPERIMENTAL PROCEDURES

### Materials

Phorbol 12-myristate 13-acetate (PMA) was from LC Laboratories (P-1680). GF 109203X (Bisindoylmaleimide I, BIM I) and AIM-100 were from Tocris-Cookson. All other reagents were from either Sigma-Aldrich or Fisher Scientific, and were of the highest possible grade.

### Animals

All studies were conducted in accordance with UMASS Medical School IACUC Protocol A-1506 (H.E.M). *Pitx3*^*IRES2-tTA*^ mice on the *C57Bl/6* background were the generous gift of Dr. Huaibin Cai (National Institute on Aging), and were continually backcrossed to *C57Bl/6* mice. Mice were maintained in 12hr light/dark cycle at constant temperature and humidity and food and water were available ad libitum.

### Antibodies

Primary antibodies used: mouse anti-Rit2 18G4 (27G2; Sigma), mouse anti-Rit2 4B5 (GTX83711, GeneTex), rat anti-DAT (MAB369), mouse anti-SERT (ST51-2; Mab Technologies), rabbit anti-HA (C29F4; Cell Signaling Technology), mouse anti-GFP (Roche), mouse anti-Ack1 (A-11; sc-28336), and rabbit anti-pY284-Ack1 (Millipore). Horseradish peroxidase-conjugated secondary antibodies: goat anti-rat, goat anti-mouse and goat anti-rabbit were from Jackson ImmunoResearch.

### Plasmids and cloning

N-S/DAT (SERT1-78/DAT60-620), DAT/C-S (DAT1-583/SERT601-630), and S/DAT/S (SERT1-78/DAT60-583/SERT601-630) plasmids, in which either the DAT N-terminus (N-S/DAT), C-terminus (DAT/C-S), or both termini (S/DAT/S) were substituted with those of SERT, were generated as previously described (53) using PCR-ligation-PCR approach to clone the DAT or SERT terminal domains onto the hSERT or hDAT-pCDNA3.1(+) backbone. CFP-tagged chimeras were generated by cloning their cDNAs into the pECFP-C1 vector using HindIII/XbaI (N-S/DAT), HindIII (DAT/C-S), and HindIII/SalI (S/DAT/S). Bungarotoxin binding site (BBS)-tagged hDAT and DAT chimera constructs were generated using the extracellular tagging strategy as previously described (49) with the following amino acid sequence inserted into extracellular loop 2 of hDAT and DAT chimera constructs: GSSGSSG<u>WRYYESSLEPYPD</u>GSSGSSG.The underlined BBS is flanked by linker sequences. All plasmids were verified by Sanger sequencing (Genewiz). Human constitutively active Ack1 mutant (S445P-Ack1-HA) was generated as previously described (24).

### AAV production and stereotaxic viral delivery

pscAAV-TRE-eGFP and pscAAV-TREmiR33-shRit2-eGFP plasmids were generated as previously described (39), and AAV particles (AAV9 serotype) were produced, purified, and titers determined by the University of Massachusetts Medical School Viral Vector Core, as previously described (70). For intracranial stereotaxic AAV injections, male and female mice (minimum 3 weeks age) were anesthetized with 100mg/kg ketamine/10mg/kg xylazine (I.P.), and 20% mannitol was administered (I.P.) 15 minutes prior to viral delivery, to increase viral spread (71). Mouse heads were shaved, placed in the stereotaxic frame, and bilateral 0.8mm holes were drilled into the skull at the indicated stereotaxic coordinates. 1µl of the indicated viruses were infused bilaterally into the VTA (Bregma: anterior/posterior: − 3.08mm, medial/lateral: ±0.5mm, dorsal/ventral: − 4.5mm) at a rate of 0.2µL/min. Syringes were left in place for a minimum of 5 minutes post-infusion prior to removal. Mice were housed for a minimum of four weeks before experiments were performed. Viral expression in each animal was confirmed by visualizing GFP expression in 0.3mm coronal ventral midbrain slices.

### Ex vivo slice biotinylation

*Ex vivo* striatal slices were prepared 4-5 weeks following viral injection in *Pitx3*^*IRES2-tTA/+*^ mice (for Rit2 KD studies), or from 5-7 week old C57Bl/6J (for PKC specificity studies). Mice were sacrificed by cervical dislocation and rapid decapitation and heads were immediately submerged in ice cold, oxygenated cutting solution, pH 7.4 (in mM: 20 HEPES, 2.5 KCl, 1.25, NaH_2_PO_4_, 30 NaHCO_3_, 25 glucose, 0.5 CaCl_2_ ·4H_2_O, 10 MgSO_4_ ·7H_2_O, 92 N-methyl-D-glucamine (NMDG), 2.0 thiourea, 5.0 Na^+^-ascorbate, 3.0 Na^+^-pyruvate) for 1 min. Brains were removed and 300µm coronal slices were prepared with a VT1200 Vibroslicer (Leica) in ice-cold, oxygenated cutting solution. Slices were hemisected along the midline, and were recovered in ACSF, pH 7.4 (in mM: 125 NaCl, 2.5 KCl, 1.24, NaH_2_PO_4_, 26 NaHCO_3_, 11 glucose, 2.4 CaCl_2_ ·4H_2_O, and 1.2 MgCl_2_ ·6H_2_O) 40 min, 31°C. Hemi-slices were treated ±1µM PMA in ACSF, 30min, 37°C with constant oxygenation, using their contralateral hemi-slice as a vehicle-treated control. Following drug treatment, slices were moved to ice and surface DAT was labeled by biotinylation as previously described (24,26,39). Briefly, slices were biotinylated with membrane-impermeant sulfo-NHS-SS-biotin (1mg/ml), 45min, 4°C. Residual biotin was quenched 2x 20 min washes of ice-cold ACSF supplemented with 100mM glycine, and were washed with ice-cold ACSF. Hemi-slices were enriched for dorsal and ventral striatum, by sub-dissecting in a line from the anterior commissure to the lateral olfactory tract. Sub-dissected slices were lysed in RIPA buffer containing protease inhibitors, and tissue was disrupted by triturating sequentially through a 200µL pipet tip, 22G and 26G tech-tips. Samples rotated 30min at 4°C to complete lysis, insoluble material was removed by centrifugation, and protein concentrations were determined using the BCA protein assay. Biotinylated proteins were isolated by batch streptavidin chromatography, overnight with rotation, 4°C, at a ratio of 20µg striatal lysate to 30µL streptavidin agarose beads, which was empirically determined to recover all biotinylated DAT. Recovered proteins were washed with RIPA buffer and eluted from beads in 2X SDS-PAGE sample buffer, 30min, room temperature with rotation. Eluted (surface) proteins and their respective lysate inputs were resolved by SDS-PAGE, and DAT was detected by immunoblotting as described above. %DAT at the cell surface was calculated by normalizing surface signals to the corresponding total DAT input signal in a given hemi-slice, detected in parallel. Note that all slice data for Rit2-KD experiments in the current study were acquired during the course of our previous study, in which we first achieved AAV-mediated Rit2 KD (39). Basal DAT surface levels in vehicle-treated male slices were compared and reported in that study, and thus were not re-analyzed for the current study. However, the DAT surface levels from vehicle-treated male hemislices were re-used in the current study as controls to determine whether PMA treatment affected DAT surface levels in contralateral hemi-slices. Rit2 knockdown in females was confirmed by RT-qPCR (Fig. S1c of previous study (39)). For males, successful viral expression (AAV9-eGFP and AAV9-eGFP-shRit2) was confirmed by visual detection of GFP reporter immunofluorescence in midbrain slices from each experimental animal. For PKC specificity studies, data were reported as %change in DAT surface levels in drug-treated hemislices, normalized to their vehicle-treated, contralateral hemi-slices.

### Cell Culture and transfection

Cells were maintained at 37°C, 5% CO2. SK-N-MC cells were grown in MEM (Sigma), and HEK293T, HEK293 (FRET studies) and N2a cells were grown in DMEM (CellGro/Corning), each supplemented with 10% fetal bovine serum, 2mM glutamine, and 100 units/mL penicillin-streptomycin. SK-N-DZ cells were grown in DMEM (ATCC #30-2002) supplemented with 10%FBS, 1X non-essential amino acids (Gibco) and 100u/mL penicillin-streptomycin. HEK293T cells were transfected using Lipofectamine 2000 (Invitrogen) according to manufacturer’s instructions with the following modifications: For biochemical and RT-qPCR studies, cells were seeded into 6-well plates at a density of 1×10^6^ (SK-N-MC), 5×10^5^ (HEK293T), or 2.5×10^5^ (N2a) cells/well one day prior to transfection, and were transfected with 3μg (SK-N-MC) or 2μg (HEK293T and N2a) plasmid DNA/well using a lipid:DNA ratio of 2:1 (SK-N-MC and HEK293T) or 4:1 (N2a). Stable cell lines were generated by selecting cells starting 48 hrs following transfection, with 0.5 mg/mL (SK-N-MC) or 0.8 mg/L (SK-N-DZ) G418 (Geneticin, Invitrogen/Life Technologies). Stably transfected cells were pooled and cell lines were maintained under selective pressure using 0.2 mg/mL or 0.32 mg/mL G418 for SK-N-MC and SK-N-DZ cells, respectively. For FRET imaging studies, HEK293 cells were seeded into an 8-well chambered coverslip (ibidi) at a density of 2×10^4^ cells/well one day prior to transfection, and were transfected with the indicated plasmids using JetPRIME (Polyplus-transfection) according to the manufacturer’s protocol. FRET studies were performed 24 hrs post-transfection.

### shRNA, Lentiviral production and cell transduction

GIPZ lentiviral shRNA constructs targeted to Rit2, and empty pGIPZ vector control, were obtained from Dharmacon. Tested shRNA clone ID’s and mature antisense sequences were as follows:

<u>shRit2-104</u>: V3LHS_380104; CTTCTTCTTCAAAGAACCT

<u>shRit2-105</u>: V3LHS_380105; TTGTTACCCACCAGCACCA

<u>shRit2-107</u>: V3LHS_380107; CTTCTTCTTCAAAGAACCT

Lentiviral particles were produced in HEK293T cells as previously described (24). For cell transduction, 1×10^6^ cells DAT-SK-N-DZ cells were seeded onto 6-well plates and were transduced with 3.0mL of lentiviral supernatant, supplemented with 0.8µg/mL polybrene, 16-24 hours post-seeding. Cells were selected for transduction beginning 24 hours post-infection (72 hrs total) with SK-N-DZ media supplemented with 1.25µg/mL puromycin. Assays were conducted 96 hrs post-transduction.

### FRET

FRET was measured using an iMIC inverted microscope (TILL Photonics GmbH). Samples were focused using a 60X (N.A. 1.49) oil objective (Olympus). Fluorescence was excited using a 100 W Xenon Lamp (Polychrome, Till Photonics GmbH). Excitation light was filtered through either a 436/20 nm (CFP) or 514/10 nm (YFP) excitation filter (Semrock) and directed to the sample via a 442/514 dual line dichroic mirror (Semrock). Emitted fluorescence light was filtered through a 480/40 nm - 570/80 nm dual emission filter (Semrock) and directed to a beamsplitter unit (Dichrotom, Till Photonics). Briefly the emission light was separated spatially according to the fluorescence wavelength using a 515 nm dichroic mirror (Semrock). The resultant two channels (<515 nm & >515 nm) were projected next to each other onto an EMCCD chip (iXon Ultra 897Andor) and recorded using Live Acquisition software (version 2.5.0.21; TILL Photonics GmbH). To guarantee the best signal to noise ratio and dynamic range, the camera was operated in 16-bit mode with a readout speed of 1 MHz. According to manufactories recommendation; an EM-Gain of 16 was applied to overcome the noise floor. To analyze FRET (see below) two images were taken per set (Donor excitation -> Donor Emission / Acceptor Emission and Acceptor Excitation -> Acceptor Emission respectively). Per condition ten sets were recorded each experimental day; the images were then analyzed using Offline Analysis software (version 2.5.0.2; TILL Photonics GmbH). One region of interest (part of the plasma membrane) per cell was selected in the CFP channel. Background fluorescence was subtracted from each image and the average intensity of each region of interest was used for calculations. Spectral bleed through (BT) for donor (0.57) and acceptor (0.04) was determined using HEK293 cells expressing a CFP or YFP signal only. Normalized FRET (NFRET) was calculated as follows:

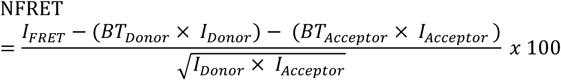

A fused CFP-YFP construct (CYFP) was included as positive control, resulting in maximum FRET. Non-fused donor and acceptor fluorophores were included as a negative control. To confirm that the calculated NFRET values reflect bona fide FRET, donor (CFP) recovery after acceptor (YFP) photobleaching (DRAP) experiments were included to support the conclusion that the fluorophore-tagged proteins directly interact at the site of the photobleaching. Average acceptor photobleaching was 85±4% (mean ±SD).

### RNA extraction & RT-qPCR

RNA was isolated from cell lines and rodent midbrain using RNAqueous®-Micro Kit RNA Isolation (Thermo Fisher Scientific). For ventral midbrain samples, 1.0 mm bilateral tissue punches, encompassing both the ventral tegmental area and substantia nigra par compacta, were taken from 300µm acute mouse and rat midbrain slices. Reverse transcription was performed using RETROscript® Reverse Transcription Kit (Thermo Fisher Scientific). Quantitative PCR was performed and analyzed using the Applied Biosystems® 7500 Real-Time PCR System Machine and Software, using Taqman® Gene Expression Assays for human Rit2 (Hs01046673_m1), Rit1 (Hs00608424_m1), and GAPDH (Hs99999905_m1), mouse Rit2 (Mm01702749_mH), and GAPDH (Mm99999915_g1), and rat Rit2 (Rn01760884_m1) and GAPDH (Rn01775763_g1).

### [^3^H]DA uptake assay

SK-N-MC cells stably expressing BBS-DAT were seeded onto 96-well plates at a density of 7.5×10^4^/well 24 hrs prior to assay. Cells were washed twice with Krebs-Ringer-HEPES buffer (120mM NaCl, 4.7mM KCl, 2.2mM CaCl_2_, 1.2mM MgSO_4_, 1.2mM KH_2_PO_4_, and 10mM HEPES, pH 7.4) and pre-incubated in KRH supplemented with 0.18% glucose ±1µM PMA, 30 min, 37°C. 100nM desipramine was included in all samples to eliminate uptake contribution of endogenous norepinephrine transporter. DA uptake was initiated by addition of 1µM [^3^H]DA (Perkin Elmer: Dihydroxyphenylethylamine (Dopamine), 3,4-[Ring-2,5,6-^3^H]) in KRH supplemented with 0.18% glucose, 10µM pargyline, and 10µM ascorbic acid. Assays proceeded for 10min, 37°C, and were terminated by three rapid washes with ice-cold KRH buffer. Cells were solubilized in scintillation fluid, and accumulated radioactivity was determined by liquid scintillation counting in a Wallac MicroBeta scintillation plate counter. Non-specific DA uptake was defined in the presence of 10μM GBR12909.

### Internalization Assays and Immunoblotting

Relative internalization rates over 10 minutes were measured by reversible biotinylation as previously described (24,25,44). Briefly, the indicated stably transfected cells were seeded into 6-well plates at 1.5×10^6^ cells/well one day prior to analysis. Cells incubated twice with 2.5 mg/mL sulfo-NHS-SS-biotin (15 min, 4°C) and quenched twice with PBS^2+^ (PBS, pH 7.4, 1.0mM MgCl_2_, 0.1mM CaCl_2_) supplemented with 100mM glycine (15 min, 4°C). Internalization was initiated by rapidly warming cells in prewarmed PBS^2+^ supplemented with 0.18% glucose, 0.2% protease-/IgG-free bovine serum albumin, and proceeded for 10min, 37°C in the presence of the indicated drugs. Parallel surface-labeled controls remained at 4°C. Cells were rapidly cooled by washing thrice with ice-cold NT buffer (150mM NaCl, 20mM Tris, pH 8.6, 1.0mM EDTA, pH 8.0, 0.2% protease-/IgG-free bovine serum albumin) and remaining surface biotin was stripped by reducing in 100mM TCEP in NT buffer twice (25 min, 4°C). Cells were rinsed rapidly in PBS^2+^, and were lysed in RIPA buffer (10mM Tris, pH 7.4, 150mM NaCl, 1.0mM EDTA, 0.1% SDS, 1% Triton-X-100, 1% sodium deoxycholate) containing protease inhibitors (1.0mM PMSF and 1.0g/mL each leupeptin, aprotinin, and pepstatin). Lysates were cleared by centrifugation and protein concentrations were determined with the BCA protein assay (Thermo Fisher) using BSA as a standard. Biotinylated proteins were recovered by streptavidin batch chromatography from equivalent amounts of cell lysate, and were eluted in 2X Laemmli sample buffer, 30 min, room temperature with rotation. Eluted proteins were resolved by SDS-PAGE and proteins were detected and quantified by immunoblotting: hDAT and DAT/C-S were detected with amino-directed rat anti-DAT (MAB369, Millipore, 1:2000), and N-S/DAT and S/DAT/S were detected with amino-directed mouse anti-hSERT (MAb Technologies,1:2000). Immunoreactive bands were detected using a VersaDoc imaging station (Bio-Rad), and were in the linear range of detection. Internalization rates were calculated as %surface protein internalized/10 min, as compared to their respective surface signal at t=0 min (controls that were biotinylated and kept at 4°C). Note that for all representative immunoblots shown throughout the study, all brightness/contrast manipulations were made uniformly across any given blot. For presentation purposes, immunoreactive bands were cropped from the same exposure of the same immunoblot.

### BBS-DAT/chimera pulldowns

HEK293T cells were transiently co-transfected with HA-Rit2, and either BBS-DAT or BBS-DAT chimeras, at a DAT:Rit2 plasmid ratio of 1:4, as described above. Cells were washed thrice with ice cold PBS^2+^ and surface BBS-DAT chimeras were labeled with 120μM biotinylated α-bungarotoxin (α-BTX-b, Thermo Fisher) in PBS^2+^, 2 hrs, 4°C. For drug treatments, cells were treated with the indicated drugs 30 min, 37°C, prior to labeling with α-BTX-b. Following labeling, α-BTX-b solution was removed, cells were washed thrice with ice cold PBS^2+^, and were lysed in ice-cold co-immunoprecipitation (co-IP) lysis buffer (50mM Tris, pH 7.4, 100mM NaCl, 1% Triton X-100, 10% glycerol, and 1.0mM EDTA) containing protease inhibitors (1.0mM PMSF and 1.0g/mL each leupeptin, aprotinin, and pepstatin) and Phosphatase Inhibitor Cocktail V (EMD Millipore), 30 min, 4°C. Labeled proteins were recovered from equivalent amounts of protein by batch affinity chromatography with streptavidin-coupled M280 Dynabeads (Thermo Fisher), 4°C, overnight with rotation. Lysate/bead ratios were empirically determined to assure quantitative recovery of all labeled proteins in lysates. Beads were gently washed thrice with ice-cold co-IP buffer, with magnetic recovery between washes, and isolated proteins were eluted from beads in an equal volume of co-IP lysis buffer and 2X SDS-PAGE sample buffer (100mM Tris, pH 6.8, 4.4% SDS, glycerol, 100mM DTT, and 0.04% bromophenol-blue). Isolated proteins were resolved by SDS-PAGE, and specific protein bands were detected by immunoblotting with antibodies for rat anti-DAT (1:2000), mouse anti-hSERT (1:2000), and rabbit anti-HA (1:2000) as indicated above. Immunoreactive HA-Rit2 band densities were normalized to their respective recovered DAT, or chimera, bands in each independent experiment.

### Statistical Analysis

All data were analyzed using GraphPad Prism software. Prior to analyses, statistical outliers within data sets were identified using either Grubb’s or Rout’s outlier test, and were removed from further analysis. Specific statistical tests used are detailed within each figure legend. For comparisons between two groups, a Student’s t test was used. For comparison among more than two experimental groups, one-way ANOVA with appropriate post-hoc multiple comparison test was performed, as indicated within each figure legend.

## Acknowledgements

We thank Tucker L. Conklin for excellent technical support. This work was supported by NIH grants DA015169 (H.E.M), DA035224 (H.E.M), F31DA039592 (C.G.S), F31DA045446 (P.J.K), Austrian Science Foundation (FWF) F35-B06 (H.H.S), and the Swiss National Science Foundation P2BSP3_181809 (D.L.). The content is solely the responsibility of the authors and does not necessarily represent the official views of the National Institutes of Health.

## Abbreviations

(DA): Dopamine
(DAT): DA transporters
(PKC): protein kinase C
(ADHD): attention-deficit/hyperactivity disorder
(ASD): autism spectrum disorder
(PD): Parkinson’s disease
(AMPH): amphetamine
(co-IP): co-immunoprecipitation
(VTA): ventral tegmental area
(SNc): substantia nigra pars compacta
(DS): dorsal striatum
(VS): ventral striatum
(BBS): bungarotoxin-binding site
(α-BTX-b): biotinylated α-bungarotoxin.

